# Probe the effect of clustering on EphA2 receptor signaling efficiency by subcellular control of ligand-receptor mobility

**DOI:** 10.1101/2021.02.16.431455

**Authors:** Zhongwen Chen, Dongmyung Oh, Kabir H. Biswas, Ronen Zaidel-Bar, Jay T. Groves

**Author notes:** Equal contribution (Zhongwen Chen, Dongmyung Oh).

## Abstract

Clustering of ligand:receptor complexes on the cell membrane is widely presumed to have functional consequences for subsequent signal transduction. However, it is experimentally challenging to selectively manipulate receptor clustering without altering other biochemical aspects of the cellular system. Here, we develop a microfabrication strategy to produce substrates displaying mobile and immobile ligands that are separated by roughly one micron and thus experience an identical cytoplasmic signaling state, enabling precision comparison of downstream signaling reactions. Applying this approach to characterize the ephrinA1:EphA2 signaling system reveals that EphA2 clustering enhances receptor phosphorylation. Single molecule imaging clearly resolves increased molecular binding dwell time at EphA2 clusters for both Grb2:SOS and NCK:NWASP signaling modules. This type of intracellular comparison enables a substantially higher degree of quantitative analysis than is possible when comparisons must be made between different cells and essentially eliminates the effects of cellular response to ligand manipulation.

## Introduction

The cell membrane surface is studded with a broad array of receptor proteins that interact with numerous ligands, which can be soluble, membrane-bound on an apposed cell surface, or associated with the extracellular matrix (Casaletto & McClatchey, 2012; Downward, 2001; Groves & Kuriyan, 2010). Ligand binding is followed by activation of elaborate signal transduction pathways in the cell, which ultimately mediate cellular decision making. In the case of receptor tyrosine kinases (RTKs), initial receptor activation after ligand binding involves phosphorylation of specific tyrosine residues on the receptors, followed by recruitment of adaptor proteins which mediate activation of additional signaling molecules. The membrane receptors, adaptors, and signaling molecules form complexes to induce immediate local responses, such as actin polymerizations, or transduce signals into nucleus, such as through mitogen-activated protein kinase (MAPK) signaling cascade. Assembly of cell surface receptors into clusters or organized arrays is a common feature of cell membranes (Dustin & Groves, 2012; Garcia-Parajo et al., 2014; Janes et al., 2012; Lee et al., 2002; Mossman et al., 2005; Salaita et al., 2010), and has long been implicated as an important factor for modulating signaling activity (Bray et al., 1998; Cebecauer et al., 2010).

More recently, receptor clustering in some cases has also been found to involve downstream signaling molecules that appear to undergo phase transitions (Banjade & Rosen, 2014; Case et al., 2019; Huang et al., 2019; Huang et al., 2017; Huang et al., 2016; Li et al., 2012; Su et al., 2016). For example, in T cell receptor signaling system, the Linker for activation of T cells (LAT) phosphorylation results in the recruitment of the adaptor protein Grb2 through binding to multiple tyrosine sites on LAT. On the other hand, Grb2 possess two SH3 domains that bind the guanine nucleotide exchange factor, Son of Sevenless (SOS), with the latter also serving as a crosslinking molecule to bridge multiple Grb2. Thus, LAT:Grb2:SOS form a 2-dimentional condensed network, or separated phase on the cell membrane (Huang et al., 2016; Su et al., 2016). Importantly, the physical form of this network determines a prolonged dwell time of SOS molecules on membrane, which ultimately establishes a kinetic proofreading mechanism to modulate the activation of SOS and its downstream effector Ras GTPase (Huang et al., 2019). In a similar fashion, the Nephrin receptor crosslinks the adaptor proteins NCK and neural Wiskott-Aldrich syndrome protein (NWASP) to form phase separated molecular assemblies on the cell membrane (Li et al., 2012). This network assembly is crucial for NWASP activation, as evident from the fact that increased NWASP dwell time in the phase-separated clusters determines subsequent actin polymerization (Case et al., 2019). These discoveries underscore the potential significance of spatial assembly of signaling molecules in modulating the chemical outcomes of the system. Furthermore, it is becoming increasingly clear that such signaling assemblies likely play an instrumental role in defining the organization of the cell membrane (Freeman et al., 2018; Kalappurakkal et al., 2019).

Although receptor assembly and phase transition processes are evident in cellular systems, it remains difficult to precisely define their functional consequences on signaling itself in living cells. A major reason for this is that chemical perturbations, such as those with pharmacological agents or mutations (Bugaj et al., 2013; Bugaj et al., 2015; Davis et al., 1994; Schaupp et al., 2014; Seiradake et al., 2013; Su et al., 2016; Wu et al., 2015), are likely to produce side effects on the cell other than modulating molecular assembly itself. For example, multi-tyrosine sites in the LAT protein mediate multivalent binding of Grb2 and SOS (Huang et al., 2016; Su et al., 2016). Reducing or increasing LAT tyrosine sites by point mutation progressively modulate clustering level and subsequent signaling responses; however, the contribution of clustering itself could not be separated from the effects of the total signaling molecule abundance and overall changes in cellular responses. In fact, this problem is universal in biophysical studies of protein organization in living cells: any perturbation that alters biological function will tend to produce a cascade of cellular effects, which can be very difficult to disentangle. An additional factor that compounds the complexity of signaling outcomes in studies involving a population of cells is the inherent cell-to-cell heterogeneity. These mandate studying individual cells in order to draw robust conclusions.

To address these issues, we seek to modulate receptor spatial organization by controlling ligand mobility, instead of perturbing intracellular components such as by creating genetic mutations or pharmacological treatments. In this respect, we have previously developed a microfabrication strategy to produce micropatterned hybrid substrates displaying ligands on both immobile, surface adsorbed polymers and mobile supported lipid membranes to simultaneously reconstitute cell-matrix and cell-cell interactions in individual cells (Chen et al., 2018). The multicomponent, micropatterned supported membrane substrates also allow for spatially segregated functionalization of different ligands to dissect crosstalk between different RTKs (K. H. Biswas et al., 2018; Kabir H. Biswas et al., 2018), in a way that is not feasible with non-patterned supported membrane substrates (Biswas, 2020; Biswas & Groves, 2019; Biswas et al., 2015; Biswas et al., 2016; Vafaei et al., 2017; Yu et al., 2015). Here we extend this method to display a ligand of interest in both mobile and immobile configurations, spatially juxtaposed on length scales small enough to enable a side-by-side comparison within an individual living cell. The immobile ligands are displayed on a functionalized poly L-Lysine-poly ethylene glycol (PLL-(*g*)-PEG) scaffold; while the mobile ligands are displayed on supported lipid membrane corrals which allows cluster formation. We employ an *in situ* UV-ozone lithographic process to selectively remove regions of the PLL-(*g*)-PEG substrate and subsequently replace these with supported lipid membranes.

Key to this strategy is that the clustered and non-clustered receptors are separated by a few microns and thus experience an identical cytoplasmic signaling state, enabling precision comparison of downstream signaling reactions. In this work, we utilize the micropatterned hybrid substrates to study the ephrinA1:EphA2 signaling system in living cells. EphA2 is a member of the Eph RTK family which is often overexpressed in aggressive breast cancers (Fox & Kandpal, 2004; Macrae et al., 2005). Clustering is thought to be integral to the ligand-mediated activation of EphA2 and downstream signaling (Davis et al., 1994; Salaita et al., 2010). When ephrinA1 is displayed on a mobile supported membrane surface, clustering upon interaction with EphA2 expressed on the surface of a living cell is readily observed. Here we reconstituted ephrinA1:EphA2 interactions in both mobile and immobile configurations in individual cells to study the effect of receptor clustering on signaling transductions. Using a series of live cell imaging and single molecule tracking experiments, we monitored local signaling events in a spatially resolved manner. The results show that clustering significantly increases EphA2 signaling efficiency by inducing greater receptor phosphorylation. Both Grb2:SOS and NCK:NWASP assemblies were observed in EphA2 clusters. Importantly, the receptor clustering consistently increases molecular binding time, which results in a modulation of their functions. Thus in accordance with other *in vitro* reconstitution studies (Case et al., 2019; Huang et al., 2019), our results provide further evidence to support kinetic proofreading mechanism of molecule activation in biomolecular condensates.

## Results

### EphrinA1 displayed on micropatterned membrane corrals are clustered by cells

With the ultimate goal of elucidating the role of receptor clustering in cellular signaling, we started by microfabricating substrates containing micron-scale supported membrane corrals displaying mobile ephrinA1 to reconstitute ephrinA1:EphA2 juxtacrine complexes on synthetic substrates (**Fig. 1A**) (Chen et al., 2018). For this, biotinylated poly-l-(lysine)-grafted-polyethylene (glycol) polymer (PLL-(*g*)-PEG-Biotin) was first coated on a glass coverslip, followed by selective deep UV lithography with a photomask (**Fig. S1A**). The deep UV-exposed PLL-(*g*)-PEG-Biotin was efficiently degraded and washed away, as ascertained by atomic force microscopy measurement after this step (**Fig. 1B**). Supported membranes were then assembled on the cleared glass area by the vesicle fusion method, and subsequently functionalized with Alexa680-labelled ephrinA1 via Ni-NTA-poly-histidine interaction (Nye & Groves, 2008). In addition, biotinylated cyclic RGD peptide was functionalized to the PLL-(*g*)-PEG-biotin through Dylight405-labelled NeutrAvidin conjugation, to engage integrins for cell spreading, thus enabling cells to interact with multiple ephrinA1-functionalized membrane (Chen et al., 2018). Fluorescent images confirmed successful functionalization of the membranes with ephrinA1, and its spatial segregation from RGD regions (**Fig. 1B**). Importantly, ephrinA1 on the supported membranes were mobile, as shown by fluorescence recovery after photobleaching (**Fig. S1B**).

**Figure 1.**
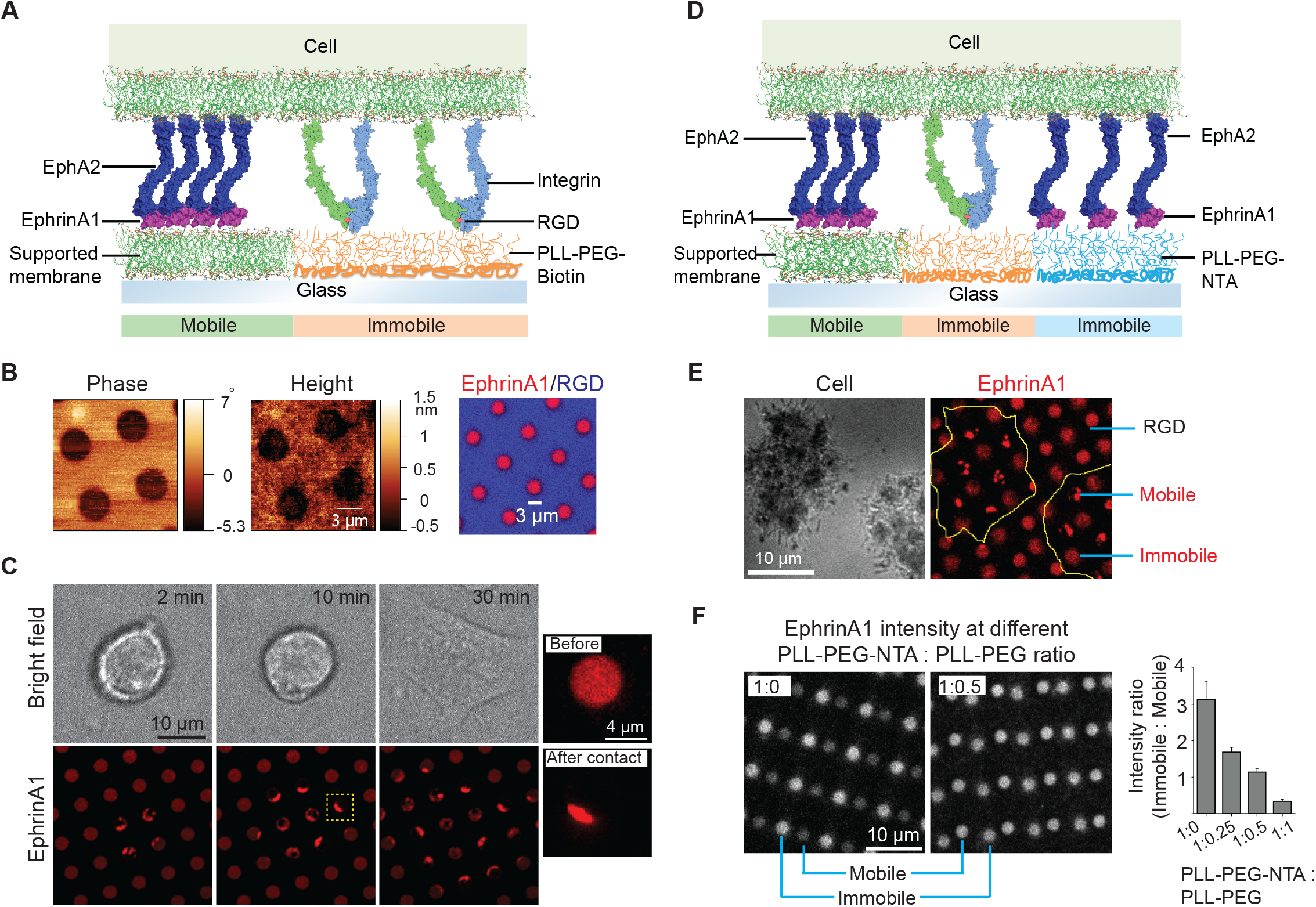
Presentation of mobile and immobile ephrinA1 ligands to a single cell. (A) Schematic illustration of 2-component substrate. (B) AFM and fluorescence images of the substrate. (C) Representative bright field and fluorescence images of a cell spreading on the substrate with indicated spreading time. Right panel represents an enlarged ephrinA1 membrane corral (marked by yellow square) before and 6 min after cell contact. (D) Schematic illustration of 3-component substrate. (E) Representative reflective interference contrast (RIC) and fluorescence images of cells spreading on the substrate. Yellow outlines indicate cell boundary. (F) Fluorescence images (in grey scale) and quantification of ephrinA1 intensity in mobile or immobile corrals by titrating PLL-(g)-PEG-NTA and PLL-(g)-PEG. Data are presented as mean ± SD.

We then monitored the interaction of live cells with ephrinA1 on the supported membrane corrals. MDA-MB-231 breast cancer cell line was used as a model cell here, since it expresses EphA2 at a high level and has been characterized previously on ephrinA1 functionalized supported membranes (Chen et al., 2018; Greene et al., 2014; Lohmuller et al., 2013; Salaita et al., 2010; Xu et al., 2011). EphrinA1 surface density on supported membranes was calibrated with quantitative fluorescence measurement (Galush et al., 2008) (**Fig. S2**), and was controlled to be around 100 molecules /μm^2^ in 4 μm-diameter membrane corrals for these experiments. Cells seeded on the micropatterned substrate spread presumably through engagement of RGD ligands on the substrate with integrins on the cell membrane. This dynamic spreading enabled cell membrane protrusions to physically interact with multiple ephrinA1 membrane corrals on the substrate, resulting in the clustering of diffusive ephrinA1 ligands through binding of cellular EphA2 receptors (Salaita et al., 2010) (**Fig. 1C** and **Movie 1**). The clustering of ephrinA1 is relatively fast: almost all the ligands inside each corral were clustered within 6 min of physical contact by the cell membrane, leading to the formation of typically a micro-cluster in each of the corrals (**Fig. 1C**). Therefore, this system allows for spatiotemporally resolved observation of ephrinA1 interaction with EphA2, as well as its downstream signaling events. The supported membranes in the micropatterned substrates could also be functionalized with other ligands, for example, E-cadherin to form cadherin junctions (**Fig. S3**), demonstrating versatility of this technology to study different receptors.

### Spatially segregated display of mobile and immobile ephrinA1 for single cells

Next, the substrates were extended to contain both mobile and immobile ephrinA1 to compare the effects of ligand clustering on receptor signaling (**Fig. 1D**). For this, PLL-(*g*)-PEG-biotin coated glass substrate was selectively UV-etched to coat another polymer PLL-(*g*)-PEG-NTA. This hybrid substrate then underwent a second UV etching for supported membranes, which led to a three-component substrate (**Fig. S4A**). The sequential UV etching processes can be overlaid randomly for regular circular arrays, because it is able to generate sufficiently large areas with clearly separated mobile and immobile regions in a centimeter-size pattern. However, the glass coverslip and the photomask can also be aligned under microscope before each UV exposure for accurate layouts of multiple components as needed, at a cost of extra time and alignment instrument (**Fig. S4B**).

EphrinA1 was functionalized on both PLL-(*g*)-PEG-NTA polymers and supported membranes through the same Ni-NTA-poly-histidine interactions, while RGD was functionalized on PLL-(*g*)-PEG-Biotin to allow cells to spread. **Fig. 1E** shows an example of cells spreading on the substrate with alternating mobile ephrinA1 membrane corrals and immobile ephrinA1 polymers. Clearly, only mobile ephrinA1 became clustered after contact with cells. Importantly, the molecular density of ephrinA1 on the polymer could be titrated by mixing PLL-(*g*)-PEG-NTA with non-reactive PLL-(*g*)-PEG (**Fig. 1F**). We controlled ephrinA1 density to be similar in both mobile and immobile regions at a range of 50-100 molecules /μ m^2^ for all cell experiments.

### Mobile ephrinA1 increases EphA2 phosphorylation through clustering

Display of mobile and immobile ephrinA1 on micron-scale corrals (membrane and polymer, respectively) allowed for comparison of clustered and non-clustered EphA2 receptor signaling in a spatially resolved manner in individual cells. Immunostaining of cells using an antibody against the intracellular domain of EphA2 verified engagement of the receptors to both mobile and immobile ephrinA1 (**Fig. 2A**). Although EphA2 cluster formation is dependent on the ligand mobility, the total amount of EphA2 receptors recruited to mobile or immobile ephrinA1 were very similar, with a slight decrease in mobile ephrinA1 area possibly due to endocytosis (Greene et al., 2014; Sugiyama et al., 2013), suggesting a conservation of binding between the receptors and ligands at the cell: substrate interface (**Fig. 2B**). EphA2 is known to undergo a ligand binding-induced autophosphorylation at tyrosine 588, and this phosphorylation site is key to the recruitment of downstream signaling molecules (Parri et al., 2005). Immunostaining of cells using an anti-pY588-EphA2 specific antibody showed that EphA2 phosphorylation increased by an average of 60% in response to mobile ephrinA1 stimulation compared to immobile stimulation (**Fig. 2A and 2C**). These results suggest that clustering of ephrinA1:EphA2 complexes enabled by the mobile ligands resulted in a change in their physicochemical properties and the level of receptor activations.

**Figure 2.**
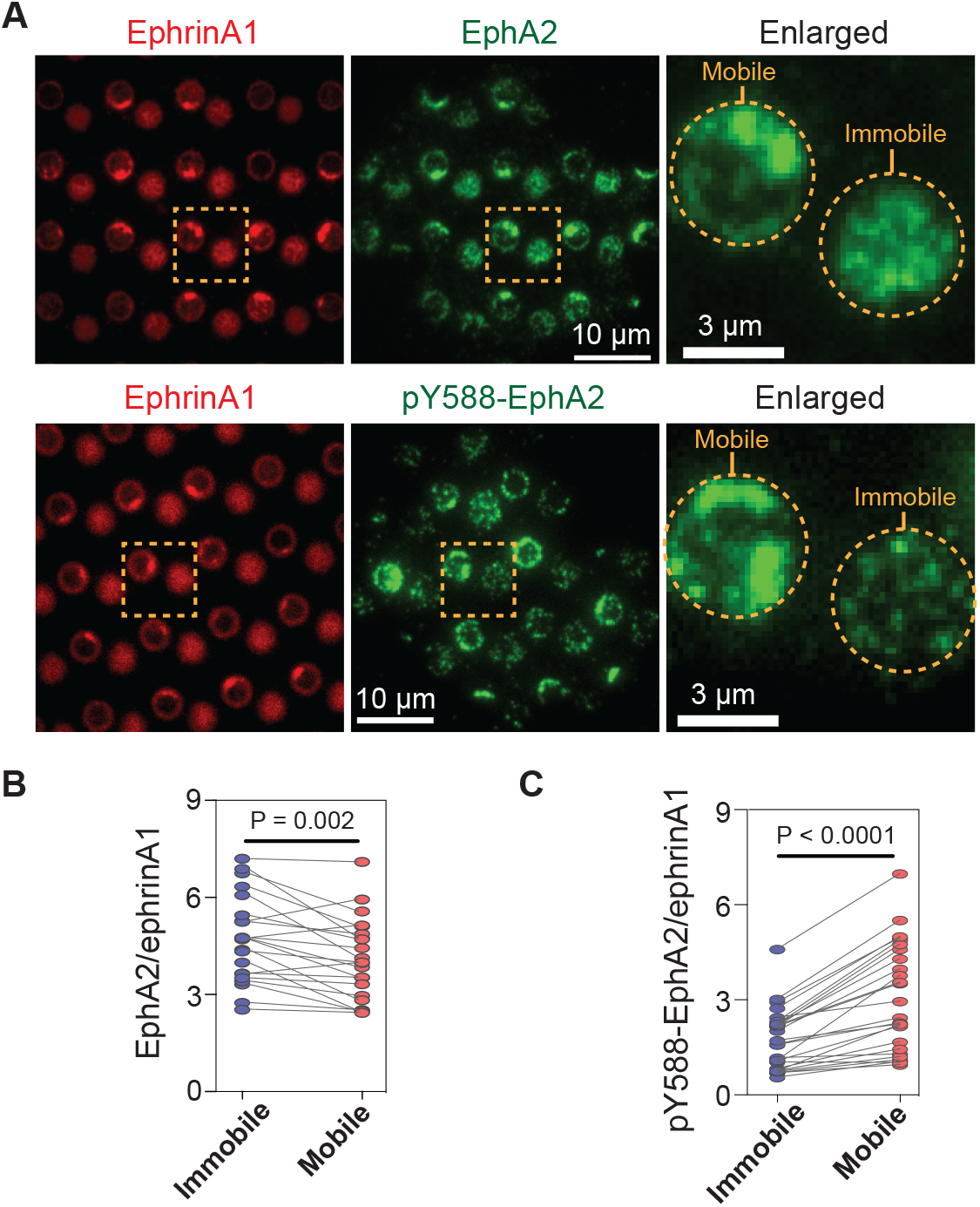
Mobile ephrinA1 stimulation increases EphA2 phosphorylation. (A) Representative immunofluorescent images of EphA2 or pY588-EphA2 in MDA-MB-231 cells fixed after 45 mins spread on the substrate. Quantification of (B) EphA2 / EphrinA1 intensity ratio or (C) pY588-EphA2 / EphrinA1 intensity ratio in mobile and immobile ephrinA1 corrals. Each data point represents an averaged ratio from multiple corrals from a single cell. The two grouped data from the same cell are paired for comparison. Significance is analyzed by paired-group student’s t test. N = 22 or 26 cells respectively.

### EphA2 clustering enhances Grb2:SOS signaling transductions by increasing on-rate and molecular dwell time

Increased phosphorylation of EphA2 observed on mobile ephrinA1 membrane corrals is expected to lead to increased recruitment of intracellular effector proteins. Therefore, we monitored the local signaling transmission events in clustered or non-clustered EphA2 receptors in single living cells. Grb2 is an important cytosolic adaptor protein that is known to be recruited to EphA2 receptors after ligand binding-induced phosphorylation (Pratt & Kinch, 2002). Grb2 further recruits SOS, which catalyzes Ras-GDP to Ras-GTP exchange, and activates the MAPK pathway (**Fig. 3A**). For this, MDA-MB-231 cells transfected with Grb2-tdEos were seeded on the substrates to allow live imaging of Grb2 recruitment by total internal reflection fluorescence (TIRF) microscopy (**Movie 2**). A remarkable difference was observed in the local recruitment of Grb2 to mobile or immobile ephrinA1:EphA2 complexes as cells spread (**Fig. 3B** and **3C**). For the same number of ephrinA1 molecules, the mobile ligands increased Grb2 recruitment by about 80% compared to immobile ones both in terms of maximal (**Fig. 3D**) and 30 mins cumulative signals (**Fig. 3E**), which is more prominent than the enhancement in receptor phosphorylation described above.

**Figure 3.**
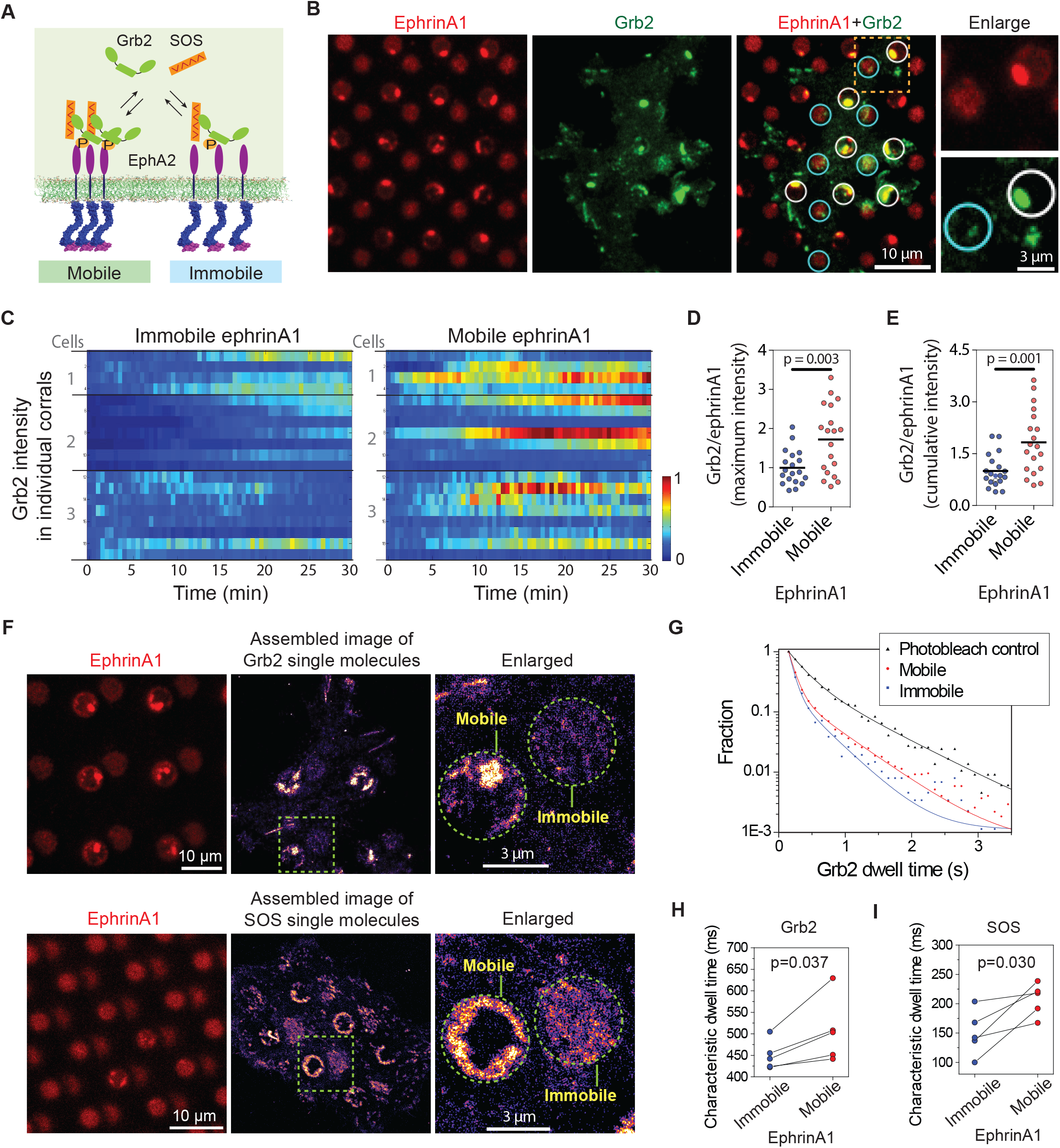
EphA2 clustering enhances Grb2:SOS signaling transductions by increasing on-rate and molecular dwell time. (A) Schematic illustration of Grb2 or SOS recruitment to mobile or immobile EphA2 receptors under the same cell. (B) Representative live cell images of a Grb2-tdEos transfected cell spreading on the substrate after 45 mins, with white circles indicating mobile ephrinA1 corrals and cyan circles indicating immobile ephrinA1. A yellow square marked region is enlarged to highlight the differential Grb2 recruitment to a mobile ephrinA1 corral and an immobile one. (C) Heat map of temporal Grb2-tdEos intensities in 3 cells. Each block represents the normalized intensity of Grb2 in an ephrinA1 corral at a given time, starting from cell contact to a total period of 30 mins, at 30 sec/frame acquisition speed. The intensity of Grb2 is normalized according to its highest intensity of all corrals through the whole time period for each cell, and color-coded for visualization. Quantification of (D) maximum Grb2 intensity or (E) 30 mins cumulative intensity in mobile or immobile ephrinA1 corrals, normalized with ephrinA1 ligand intensity in each corral. Significance is analyzed by student’s t test. (F) Single molecule imaging of Grb2-tdEos or SOS-tdEos. The coordinates of Grb2 or SOS single molecules when they first appear in a continuous movie are assembled to generate a localization image. (G) Distribution of Grb2-tdEos single molecule dwell time. Membrane located CAAX-tdEos is applied as a photobleaching control measured from another cell. Quantification of (H) Grb2-tdEos or (I) single molecule dwell time. The dwell time distribution is fitted by a 2-order exponential decay function and the slower time constant τ2 is used to represent characteristic dwell time for pairwise comparison in a group of cells. N = 5 cells. Significance is analyzed by paired-group student’s t test.

Grb2 and SOS assembly increases their molecular dwell time on the membrane, which forms a gating mechanism to control SOS activation in a reconstitution system (Huang et al., 2019; Huang et al., 2016). In our living cell system, the simultaneous presentation of clustered and non-clustered EphA2 in a single cell enables a precise comparison of Grb2 dwell time (or off-rate, K_off_ = 1/dwell time) by single molecule imaging. Grb2-tdEos fluorescence emission spectrum can be photo-switched by UV excitation. A short UV (405nm) exposure resulted in the illumination of a small amount of Grb2-tdEos in the red channel, which were then imaged at a rate of 20 frames per second (**Movie 3**). The Grb2-tdEos particles were confirmed to be single molecules as shown by single-step photobleaching, and the fluorescence intensities of individual particles also exhibited a unimodal distribution (**Fig. S5**). The coordinates of every Grb2 molecule in a continuous movie were then assembled to generate a high-resolution localization image (also termed single particle tracking Photo-Activation Localization Microscopy; sptPALM (Manley et al., 2008)), which clearly showed clustering of Grb2 in mobile ephrinA1 corrals while relatively uniformly distributed in immobile corrals (**Fig. 3F**). By mask-separating these two regions, we found that Grb2 has a longer dwell time distribution on clustered ephrinA1-EphA2 complexes, in comparison to the non-clustered ones (**Fig. 3G**). A membrane localized CAAX-tdEos control was used to measure the photobleaching rate under the same experimental condition, which was significantly slower than the apparent Grb2 binding dynamics. The dwell time difference is consistent in all measured cells as shown by pairwise comparison of mobile or immobile regions in each individual cell (**Fig. 3H**). Notably the cell-to-cell variations of dwell time are in the similar level to the mobile-to-immobile differences; and therefore, only side-by-side comparison of each individual cell made it possible to detect the increase of Grb2 membrane dwell time induced by clustering.

Similar to the observations with Grb2, EphA2 clustering also consistently increases SOS-tdEos single molecule dwell time on the membrane (**Fig. 3F and 3I**). Taken together, these data indicate that EphA2 clustering not only increases Grb2 and SOS abundance by inducing higher receptor phosphorylation, but also increases their dwell time, all of which contributing to an enhanced signaling transduction from the ephrinA1:EphA2 complexes.

### EphA2 clustering enhances NCK:NWASP induced actin polymerizations and increases molecular dwell time

Actin polymerization is another important signaling module that has been studied extensively (Banjade & Rosen, 2014; Case et al., 2019; Li et al., 2012). It was found that in a reconstitution system, receptor phosphorylation dependent NCK:NWASP assembly facilitates Arp2/3 complex activation and actin polymerizations, in which the stoichiometry regulated NWASP dwell time determines the strength of signaling responses (Case et al., 2019). EphA2 has also been reported to modulate actin cytoskeleton (Mohamed et al., 2012). Here we sought to test if such molecular timing mechanism applies to EphA2 signaling in living cells (**Fig. 4A**).

**Figure 4.**
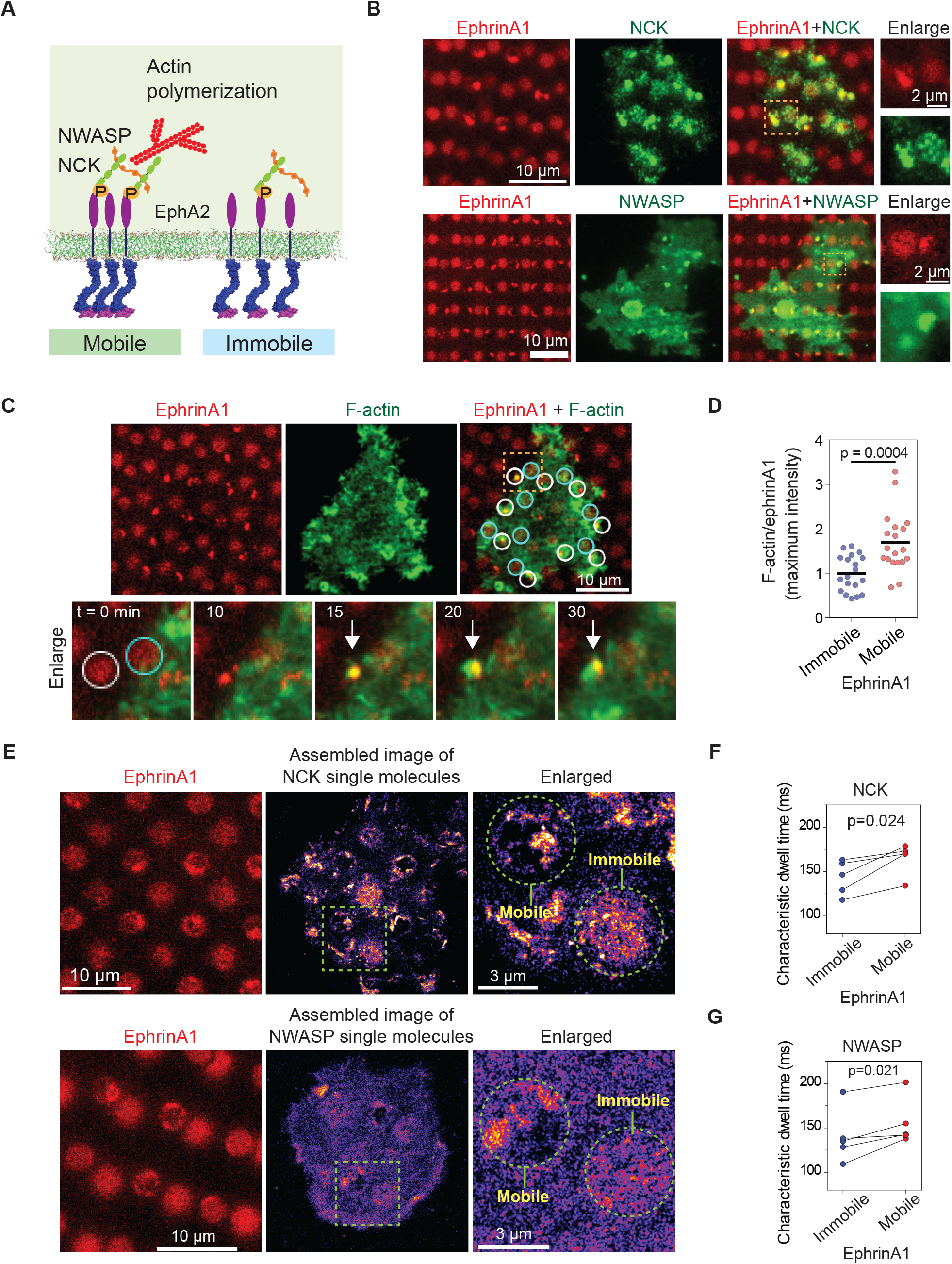
EphA2 clustering enhances NCK:NWASP induced actin polymerizations and increases molecular dwell time. (A) Schematic illustration of the signaling pathway from EphA2 receptor to actin polymerization. (B)Representative live cell images of a NCK-mEOS3.2 or NWASP-mEOS3.2 transfected cell spreading on the substrate after 1 hour. (C) Representative live cell images of a F-tractin-EGFP transfected cell spreading on the substrate, with white circles indicating mobile ephrinA1 corrals and cyan circles indicating immobile ephrinA1. The marked yellow square is enlarged to highlight temporal actin polymerization dynamics with referring to mobile or immobile ephrinA1 contact. The white arrow indicates local actin polymerizations. (D) Quantification of maximum F-trac-tin-EGFP intensity in mobile or immobile ephrinA1 corrals, normalized with ephrinA1 ligand intensity in each corral. Significance is analyzed by student’s t test. (E-G) Single molecule imaging and quantification of NCK-mEos3.2 and NWASP-mEos3.2. Condition same as Fig. 3 (F-I). N = 5 cells. Significance is analyzed by paired-group student’s t test.

Similar to Grb2, NCK-mEOS3.2 and NWASP-mEOS3.2 were clearly found to be enriched at ephrinA1:EphA2 clusters in comparison to immobile ephrinA1 region (**Fig. 4B**). Live imaging of F-tractin-EGFP showed local actin polymerization in each mobile ephrinA1 corrals shortly after cluster formation; however, no enrichment of F-actin can be resolved in regions of immobile ephrinA1 (**Fig. 4C** and **movie 4**). The average maximal intensity of F-tractin-EGFP increased by about 80% in EphA2 clusters compared with immobile corrals (**Fig. 4D**), showing that only clustered EphA2 are effective to trigger signaling transduction towards actin polymerizations. Single molecule imaging confirmed clustering of NCK-mEos3.2 and NWASP-mEos3.2 in mobile ephrinA1 corrals (**Fig. 4E**). The molecular dwell time also increased consistently in all measured cells in EphA2 clusters (**Fig. 4F and 4G**). Therefore the live cell measurement is in agreement with in vitro reconstitution experiments (Case et al., 2019), that the increased dwell time of NWASP by crosslinking is important for its activation.

### EphA2 clustering increases Grb2 and NWASP dwell time in COS7 cells

The model cell line MDA-MB-231 expresses a very high level of EphA2 receptors, which may raise concern whether the observed dwell time difference is due to EphA2 pre-clustering. We validated these results by testing another cell line COS7, which expresses EphA2 at a low to moderate level (Sabet et al., 2015). EphrinA1 clustering and subsequent cytosolic Grb2 and NWASP recruitment upon cell contact is readily observed in mobile membrane corrals (**Fig. 5A**). The clustering consistently increases Grb2-tdEos dwell time in all measured cells (**Fig. 5B**), and increases NWASP-mEOS3.2 dwell time in most cells only with a rare exception (**Fig. 5C**).

**Figure 5.**
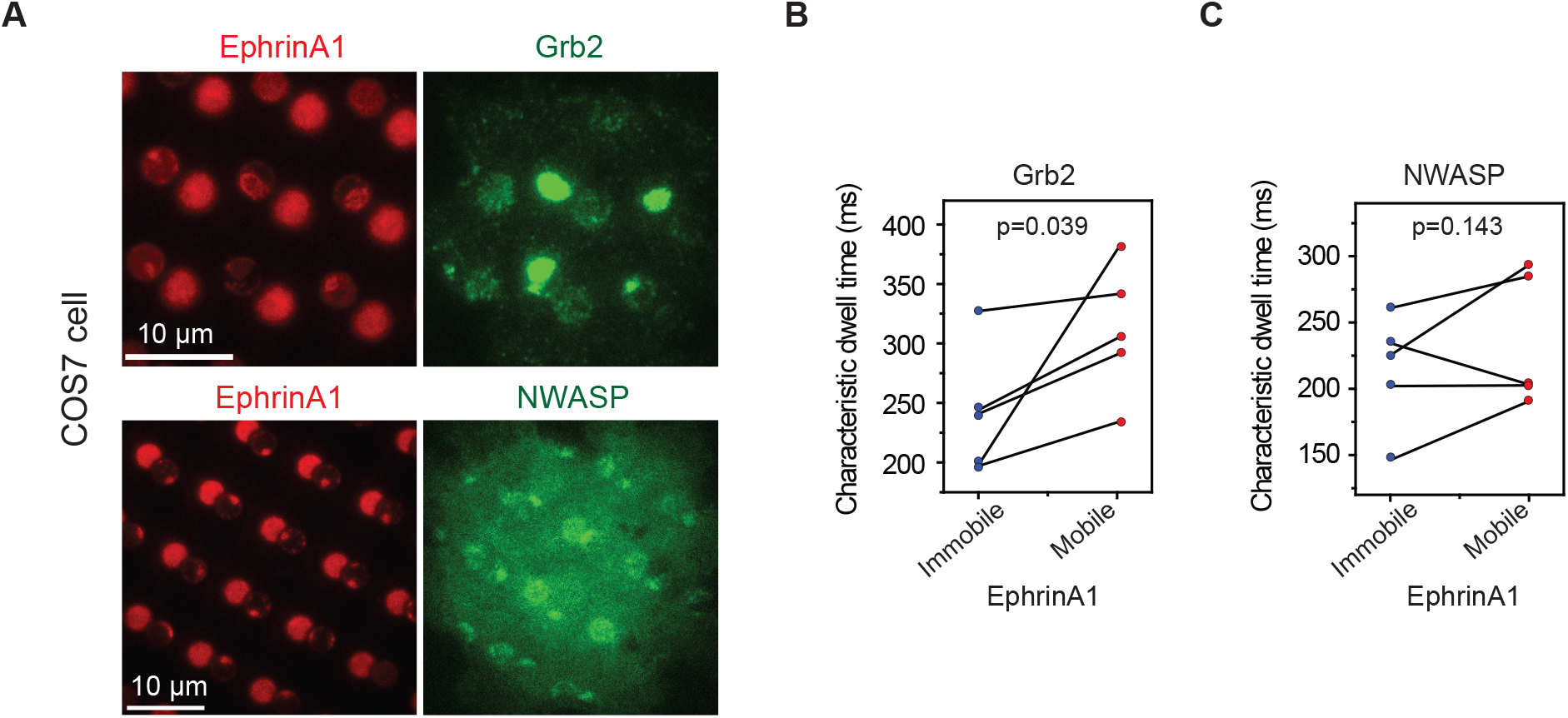
EphA2 clustering increases Grb2 and NWASP dwell time in COS7 cells. (A) Representative live cell images of Grb2-tdEOS or NWASP-mEOS3.2 transfected COS7 cells spreading on the hybrid substrate. Quantification of (B) Grb2-tdEos or (C) NWASP-mEos3.2 dwell time in COS7 cells. Condition same as Fig. 3 (F-I). N = 5 cells. Significance is analyzed by paired-group student’s t test.

## Discussion

Receptor clustering lies at the center of incorporating extracellular signals into amplified cellular responses (Bray et al., 1998; Cebecauer et al., 2010; Taylor et al., 2017). In this respect, lateral diffusion of membrane proteins is critical in controlling their clustering and thus the assembly of signaling complexes. The microfabrication technology described here provides a straightforward way to produce synthetic substrates displaying mobile as well as immobile ligands for monitoring clustering-dependent receptor signaling in individual cells. The utility of this technology is clearly demonstrated by the observation of increased phosphorylation of clustered EphA2 compared to the non-clustered ones. Our results indicate that ephrinA1 ligand binding is not sufficient to induce EphA2 receptor autophosphorylation. Instead, the increased local density of ephrinA1: EphA2 complexes as seen on mobile membrane corrals is likely to increase the accessibility of receptor catalytic sites, which are coupled with their allosteric activation (Wiesner et al., 2006). It is also possible that phosphatases are physically excluded from central kinase-dominated core to maintain EphA2 phosphorylation in the clusters, in a manner similar to that observed in the immunological receptors (Davis & van der Merwe, 2006; James & Vale, 2012). Possibly a combination of these mechanisms gives rise to the amplified EphA2 activation when they are clustered.

Two recent studies proposed a kinetic proofreading model in biomolecular condensates (Case et al., 2019; Huang et al., 2019). The key principle is that there is a time-lag between molecular recruitment and activation. Therefore, molecular timing functions as an important gating mechanism to drive nonequilibrium signaling responses: the long-dwelled molecules in phase-separated clusters are likely to be more productive in multi-step enzymatic reactions. This model was primarily inferred from simplified in vitro reconstitution experiments. Here we found that both Grb2:SOS and NCK:NWASP signaling molecules are condensed in EphA2 clusters in living cells, possibly forming a 2-dimensional network. Importantly, these molecules dwelled longer inside the clusters, which is in agreement with kinetic proofreading model. Only side-by-side comparison of mobile and immobile receptors made it possible to detect such differences in living cells.

However, the live cell results still differ from reconstitutions to some extent: the increase of molecular dwell time by receptor clustering is much smaller than it was predicted from in vitro reconstitution model, which can be several folds longer. There are several possible reasons for this. 1) Signaling transduction is far more complex in live cells. The molecular assembly is constantly modulated by phosphatases and competitive binding partners, which also change over time. 2) There possibly has only a very small fraction of super-long dwelled molecules that function most significantly in signaling transductions, as proposed in the reconstitution model (Huang et al., 2019; Huang et al., 2016). Such a small population made it very difficult to be detected in living cells. 3) Single molecule imaging is unavoidably affected by photobleaching. The absolute dwell time is almost impossible to be deconvolved from photobleaching effect in current live cell measurements, given the fact that the optical performance of fluorescent proteins cannot match organic dyes used in vitro. Higher spatial and temporal resolution are still needed to delineate more detailed mechanisms how biomolecular condensation modulate signaling efficiency.

In conclusion, EphA2 clustering enhances both receptor phosphorylation and binding dwell time of downstream signaling molecules in living cells, providing further evidence to support the kinetic proofreading mechanism of molecular activation in biomolecular condensates. While we have investigated EphA2 receptors in the current study, the technology described here could be extended to investigate other membrane-localized receptor systems. Further, the substrates developed here could be directly utilized in the screening of pharmacological agents to target diseases that involve clustering-dependent receptor signaling (Chang et al., 2018; Lohmuller et al., 2013; Salaita et al., 2010).

## Materials and Methods

### Patterned substrate of mobile and immobile ligands

Phospholipid vesicles were prepared using previously published methods (Galush et al., 2008). 96% of DOPC (1,2-dioleoyl-sn-glycero-3-phosphocholine) was mixed with 4% of Ni-NTA-DOGS (1,2-dioleoylsn-glycero-3-[(N-(5-amino-1-carboxypentyl)iminodiacetic acid)-succinyl]) (Avanti lipids) to form supported membranes. The pattern is fabricated by UV lithography. Briefly, clean coverslips were incubated with PLL-(*g*)-PEG-biotin (50%, Susos) for at least two hours. The polymer coated coverslips were then etched by deep UV (UVO-cleaner 342-220, Jelight) with a designed photomask to generate patterns. Lipid vesicles were then deposited on the coverslips to form supported membranes on the etched surface only. Later, the substrates were blocked by 0.05% bovine serum albumin (BSA; Sigma-Aldrich) for 2 hours, and then incubated with 1 μ g/ml of DyLight-405 NeutrAvidin (Thermo-Fisher) for 30 minutes to bind biotin. After rinse, the substrates were then incubated with a solution of 5 nM of ephrinA1-Alexa 680 and 1 μ g/ml of RGD-PEG-PEG-biotin (Peptides international) for 60 min for surface functionalization. EphrinA1 was expressed with a C-terminal 10-histidine tag and purified from insect cell culture (Xu et al., 2011), and labelled with Alexa 680 fluorophore (Thermo-Fisher) followed by vender’s manual. Finally, the protein incubation solutions were exchanged with imaging buffer (25 mM Tris, 140 mM NaCl, 3 mM KCl, 2 mM CaCl_2_, 1 mM MgCl_2_, and 5.5 mM D-glucose) and warmed up to 37 °C prior addition of cells.

The three component substrates were prepared by two steps UV etching. After the first etch on PLL-(*g*)-PEG-biotin surface, the coverslip was then coated with PLL-(*g*)-PEG-NTA or PLL-*g*-PEG (Susos), and underwent another etch to form supported membranes. The two etch processes were overlaid randomly for regularly circular patterns, or aligned under microscope to obtain high accuracy. The substrate was then blocked by BSA and functionalized with ligands similar as above. To modulate immobile / mobile ephrinA1 intensity ratio, PLL-(*g*)-PEG-NTA was titrated by mixing with PLL-(*g*)-PEG. The fluorescence intensity of ephrinA1 is measured 30 mins after washing out proteins in solution, at a similar time when the cellular experiments are performed.

### Cell culture, plasmid, and immunostaining

MDAMB231 cells (ATCC) and COS7 cells (UC Berkeley cell culture core facility) were grown in DMEM (high glucose) (Thermo-Fisher) and RPMI 1640 (Thermo-Fisher) medium respectively, supplemented with 10% fetal bovine serum (FBS) (Thermo-Fisher) and 1% penicillin/streptomycin (Thermo-Fisher), and kept at 37 °C in an atmosphere of 5% CO2. Cells were detached by enzyme free dissociation buffer (Thermo-Fisher) and then allowed to interact with indicated substrates. Cells were transfected by Lipofectamine 2000 (Thermo-Fisher). Grb2-tdEos, SOS-tdEos and CAAX-tdEos plasmids are the same as previously published (Oh et al., 2012). F-tractin-EGFP is maintained in Zaidel-Bar’s lab. NCK-mEos3.2 and NWASP-mEos3.2 are cloned in Mechanobiology Institute Core facility. For immunostaining, cells were fixed with 4% paraformaldehyde, and permeabilized with 0.1% Triton-X, followed with standard immunostaining protocol. Primary antibodies include rabbit anti EphA2 (CST, 7997) and rabbit anti pY588-EphA2 (CST, 12677). The secondary antibody is goat anti rabbit antibody conjugated with Alexa 488 fluorophores (Thermo-fisher).

### Live cell and single molecule imaging

An Eclipse Ti inverted microscope (Nikon) with a TIRF system and Evolve EMCCD camera (Photometrics) was used for live cell imaging. TIRF microscopy was performed with a 100x TIRF objective with a numerical aperture of 1.49 (Nikon) and an iChrome MLE-L multilaser engine as a laser source (Toptica Photonics). Immunofluorescent imaging was also acquired in an Eclipse Ti inverted microscope (Nikon) with CSU-X1 confocal spinning disk unit (Yokogawa).

Time-lapse single molecule imaging of Grb2-tdEos, SOS-tdEos, NCK-mEos3.2 and NWASP-mEos3.2 were performed by TIRF microscopy, in a way such as to optimize signal-to-noise and temporal resolution by coupling minimizing laser power and maximizing video rate. To increase tracking accuracy, the density of individual molecules was controlled by 405 nm laser illumination to be about ~0.5 / μ m^2^. Far-red channel (ex =647 nm, em > 655 nm) were acquired before single molecule recording to localize mobile and immobile ephrinA1 corrals. The autofluorescence on the red channel was completely photobleached before photo-switching Eos by a 405 nm beam. After photo-switching, a small amount of Eos molecules were visualized and recorded by EMCCD with 20 frame per second video rate. Each movie contains 1000 frames for further analysis. Membrane localized CAAX-tdEos movies were used to calculate photobleaching rate, acquired at the same microscopic set up.

### Image analysis

Live cell and immunofluorescence images were analyzed to quantify Grb2-tdEos, F-tractin-EGFP, anti-EphA2, and anti-pY588-EphA2 intensities in mobile and immobile ephrinA1 regions. The regions of mobile and immobile ephrinA1 were outlined to generate masks, so the average intensity of different channels can be measured in the same corral and the ratios were calculated after subtraction of noises.

For live cell single particle tracking, a cross-correlation single particle tracking method was used to determine the centroid positions of tdEos or mEos3.2 single molecules (Oh et al., 2012; Oh et al., 2014). A trajectory was created by connecting the subsequent xy coordinates through the frames using the nearest neighbor method. The first positions of each trajectory were assembled to generate a localization image. By using pre-acquired ephrinA1 image as a mask, single molecule signals coming from mobile or immobile ephrinA1 regions were separated to calculate spatially resolved binding kinetics. The dwell time distributions of molecules in the two different regions were fitted with a 2-order exponential decay function y=y0+A1e^−t/τ1^+ A2e^−t/τ2^, providing two characteristic time constants τ1 and τ2. To be consistent τ2 were used for pairwise comparison of each cells.

## Acknowledgement

We thank Professor Michael Sheetz for stimulating discussions. This work was supported by National Institutes of Health, National Cancer Institute Physical Sciences in Oncology Network Project 1-U01CA202241. Collaborative work at the Mechanobiology Institute, National University of Singapore, was supported by CRP001-084. Z.C is also funded by Shanghai Municipal Science and Technology Major Project ZJLab (2018SHZDZX01)

## Supplementary information for

**Fig. S1.**
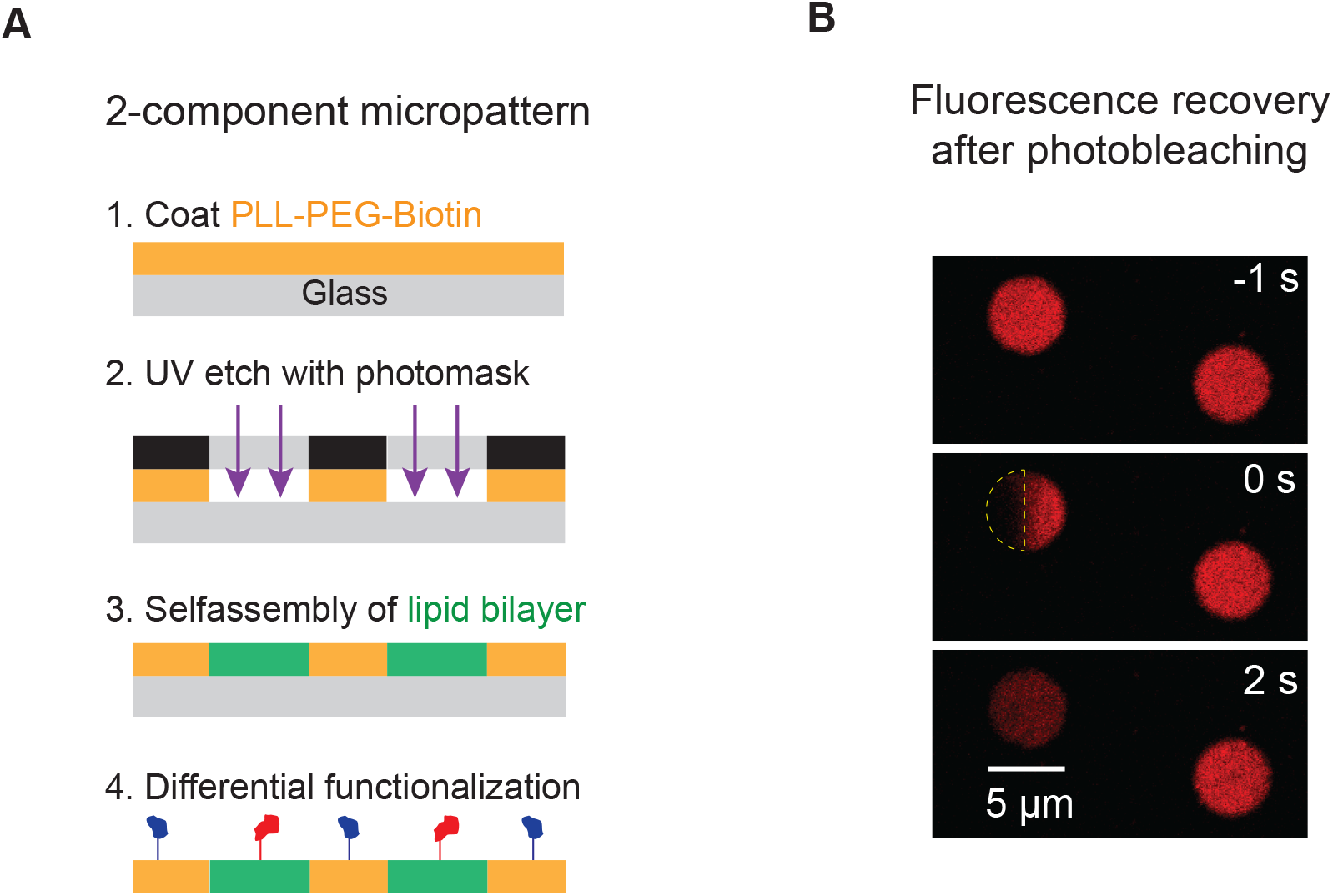
2-component patterned substrate. (**A**) Schematic illustration of the patterning method. (**B**) FRAP images of ephrinA1 in a micropatterned membrane corral, with indicated time before and after photobleach. Yellow semicircle indicates the photobleach area. The recovery of fluorescence signal in the photobleached area with a concomitant decrease in the total fluorescence intensity of the corral indicates effective ‘fencing’ of the membranes.

**Fig. S2.**
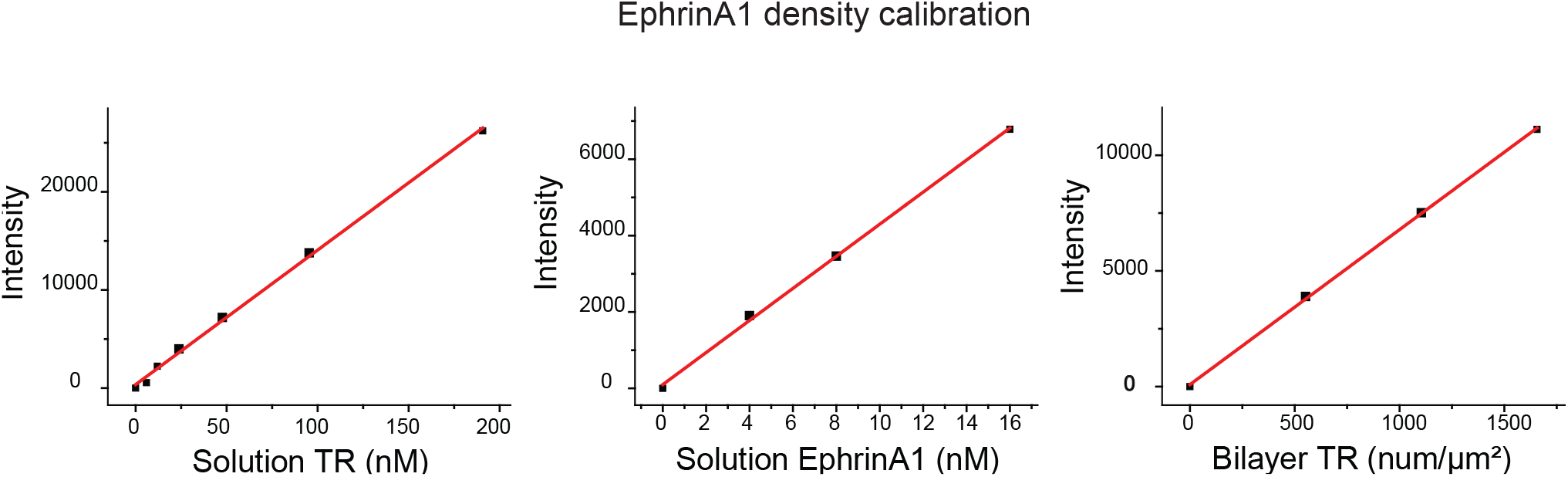
Calibration of ephrinA1 density on supported membranes. EphrinA1 density on lipid bilayers was calibrated by comparing fluorescence intensity ratio in solution and on bilayer with a reference marker Texas Red (TR). At a given imaging seting, F = constant = I(EphrinA1_solution_) / I(EphrinA1_bilayer_)= I (TR_solutio_n) / I(TR_bilayer_), where I is the concentration-normalized intensity of fluorophores. I(TR_solution_), I(EphrinA1_solution_), and I(TR_bilayer_) are calculated from linear fiting of the data points in the three graphs. Therefore, I(EphrinA1_bilayer_) = I(EphrinA1solution) x I(TR_bilayer_) / I (TR_solution_) = 20.52 intensity(a.u.) / density (num/μ m^2^). The final density of EphrinA1 on bilayer is easily calculated as (measured fluorescence intensity) / 20.52 num/μ m^2^.

**Fig. S3.**
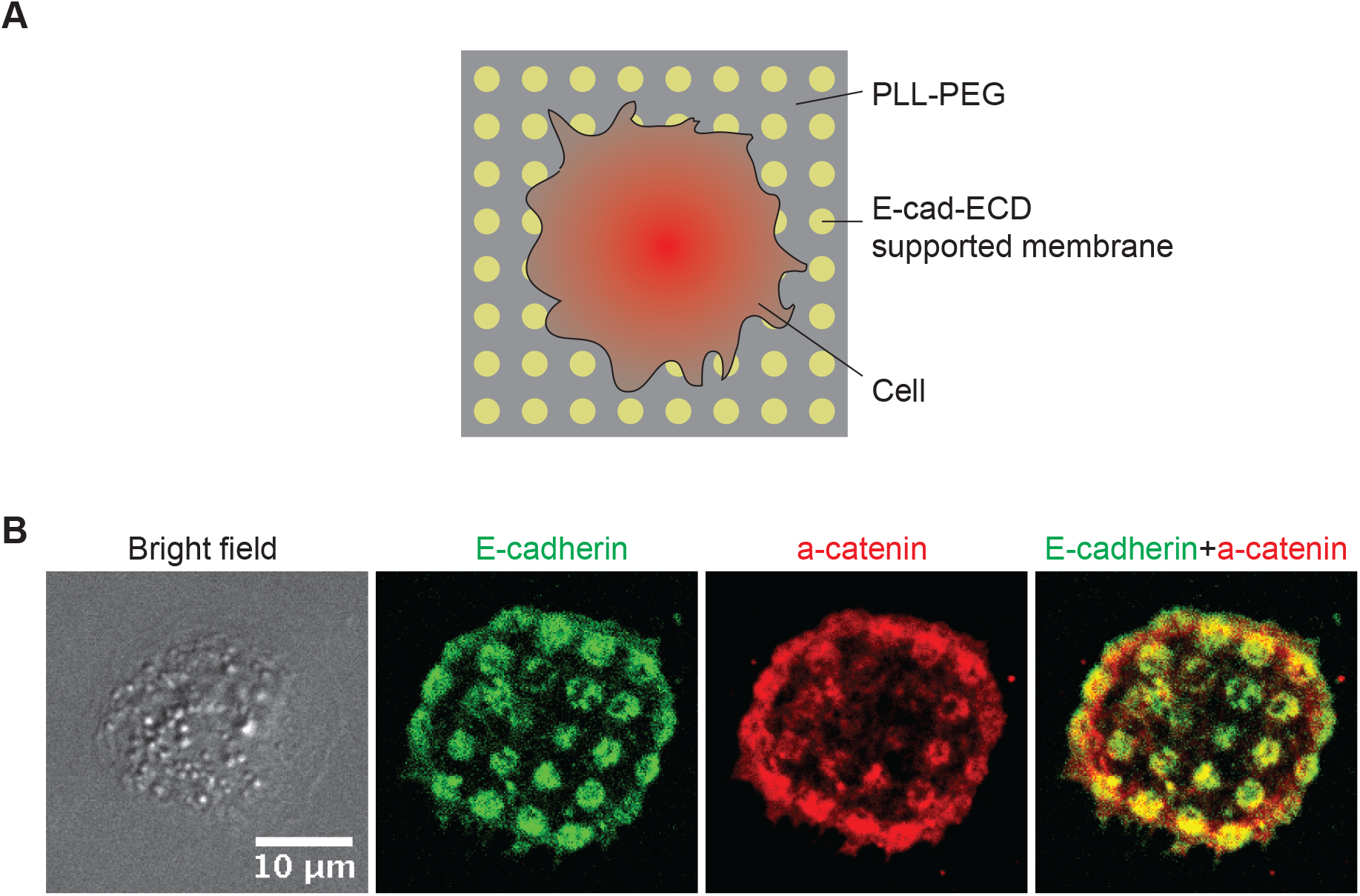
Reconstitution of E-cadherin adhesion on micropatterned supported membrane substrates. (A) Schematic illustration of cells adhering to a micropatterned substrate containing E-cadherin extracellular domain (E-cad-ECD) functionalized supported membrane corals. (B) Bright field and fluorescent images of E-cadherin-GFP and α-catenin showingthe association of α-catenin to E-cadherin clusters.

**Fig. S4.**
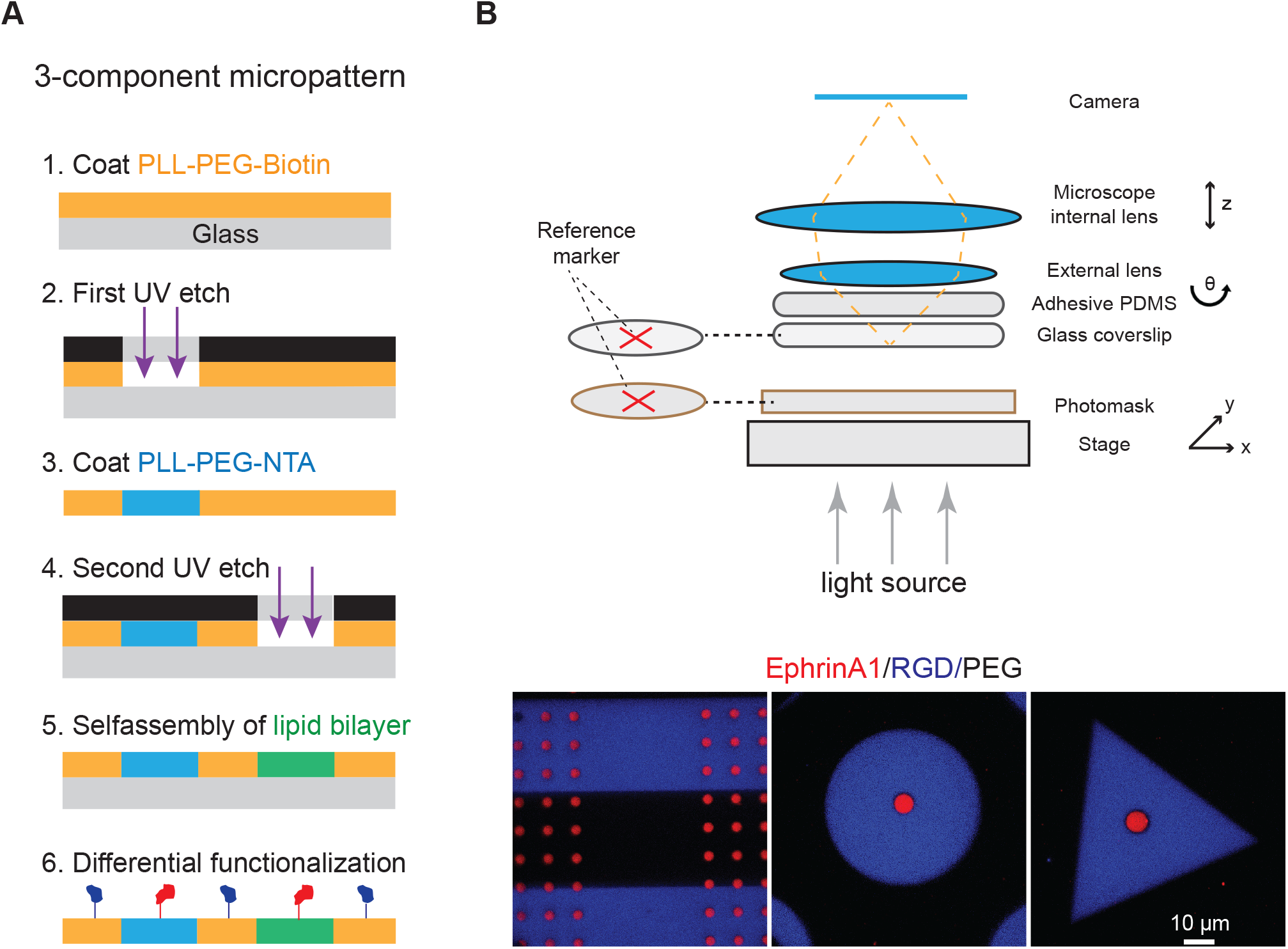
3-component patterned substrate. (**A**) Schematic illustration of the patterning method. (**B**) Versatile layouts of the three-component substrate by microscopy assisted alignment. Before each UV etch process, the maker on the polymer coated coverslip and the one on the photomask are aligned at the same position so that the three components layout can be controlled as design.

**Fig. S5.**
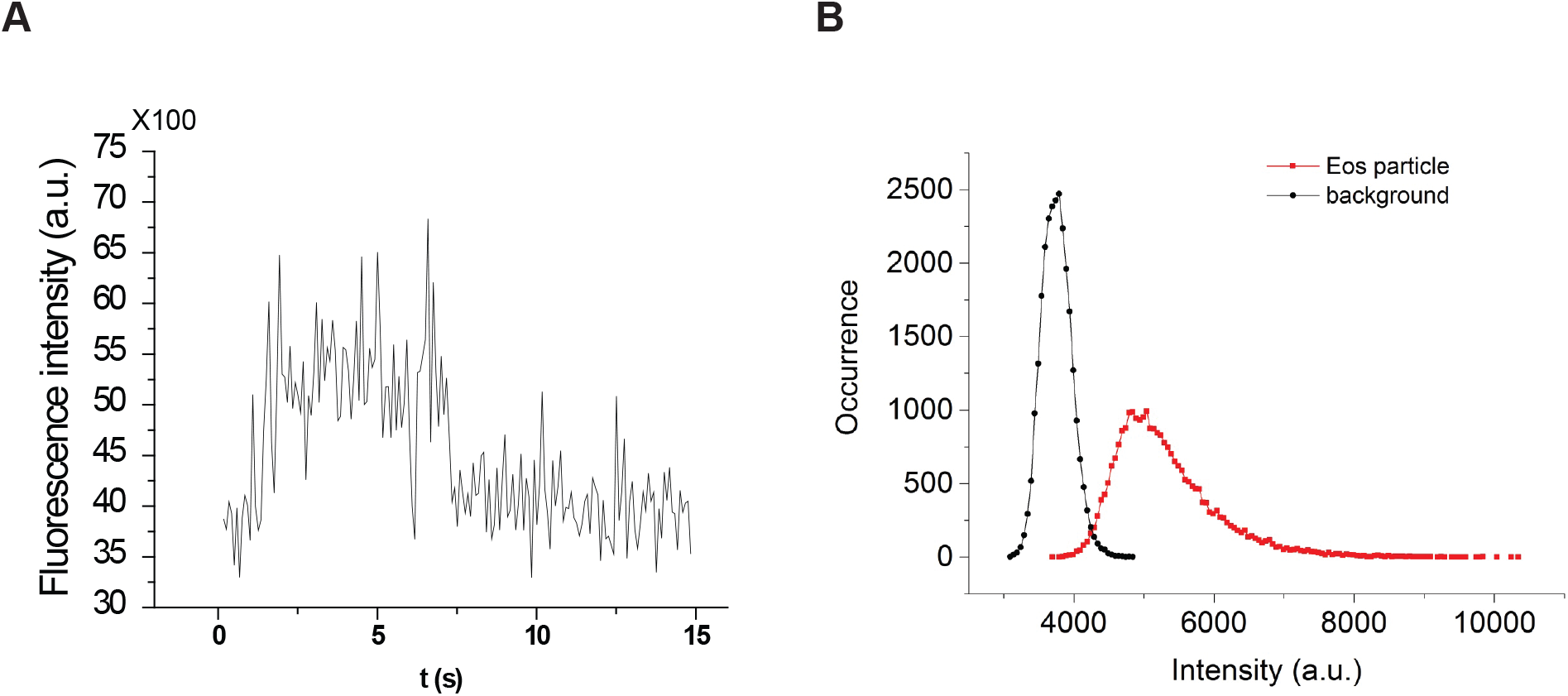
Grb2-tdEos single molecule imaging characterization. (**A**) Single step photobleach of Grb2-tdEos in live cells. (**B**) Distribution of Grb2-tdEos fluorescence intensities in a live cell movie.

